# The information content of species: formal definitions of pangenome complexity track with bacterial lifestyle

**DOI:** 10.1101/2025.03.28.645969

**Authors:** Apurva Narechania, Dean Bobo, M Thomas P Gilbert, Shyam Gopalakrishnan

**Affiliations:** Institute for Comparative Genomics, American Museum of Natural History, New York, NY, USA; Center for Evolutionary Hologenomics, The Globe Institute, University of Copenhagen, Copenhagen, Denmark; Department of Ecology, Evolution, and Environmental Biology, Columbia University, New York, NY, USA; University Museum, NTNU, Trondheim, Norway

## Abstract

Genes and other genomic elements have variable presence absence patterns across most bacterial species. Pangenome fluidity is often invoked to measure this genome flux. Fluid pangenomes contain genes found only in subsets of species strains. Tighter pangenomes contain more genes that define a shared core. Species definitions are often tied to this pangenome diversity. In any global comparative framework, pangenomes must be calculated across all known species. But defining pangenomes is fraught with computational and biological challenges, requiring assembly, annotation, alignment, and phylogenetics of millions of orthologs. We offer an alternate view that de-centers the gene and emphasizes the raw information content of sequences. Information is data that reduces uncertainty. Tight pangenomes, with elements repeated across every strain in a species ensemble, contain more complete information. In contrast, fluid pangenomes have more uncertainty, higher complexity, and higher information diversity. Bacterial lifestyle has been shown to drive this information diversity. For example, challenging environments often increase information diversity by encouraging the accrual of auxiliary genes. Here, we employ agile complexity metrics to quantify this increase. Ensembles of free-living, motile, and non-pathogenic species have high genomic complexity. Ensemble complexity decreases in species bound to specific hosts. Because we eliminate annotation and alignment, our method is fast enough to evaluate species across all known bacterial genomes. The approach democratizes classification and our results highlight how broad the term “species” has become.

## Introduction

Species definitions are starkly different across the microbial tree of life^1^. Microbes come in a variety of flavors. Some species are composed of a few very similar strains, while others have hundreds of strains whose genomes vary substantially in gene content. Pangenomes summarize this variation by outlining a set of core genes shared across strains, and a set of accessory genes found across various subsets. Pangenome size and complexity can therefore vary considerably^2,3^.

Pangenome complexity was first defined as the rate of gene accumulation with whole genomes sequenced^4^. Researchers will typically sequence a set of strains and assemble, align and annotate them. Tree building and orthology estimation follow. This set of orthologs is then split into a core/shell dichotomy, and metrics like pangenome fluidity quantify their relative weights^5^. A tighter core implies more shared genes, a less fluid pangenome, and a pangenome curve that saturates. Alternatively, genes may be found sporadically across strains, and the curve may never reach a clear asymptote.

Pangenome complexity has been shown to vary with bacterial lifestyle^6^. For example, *Prochlorococcus marinus* has 60,000 genes in its pangenome many of which are restricted to subsets of strains^7^. *Mycobacterium tuberculosis* has a mere 3500, most shared across all strains in the species^8^. Clearly, these organisms employ very different evolutionary strategies suited to their environmental niches. *P. marinus* is a free-living marine microbe that must contend with the open ocean. It lives in a milieu of candidate genes for horizontal transfer. Its large pangenome could be interpreted as a library of diverse information necessary to survive and flourish in its harsh setting, or the information capacity required to compute its environment. *M. tuberculosis* is instead restricted to the lungs of its host. Its small pangenome requires only the set of genes necessary to flourish in a uniform environment. *P. marinus* and *M. tuberculosis* are at opposite poles on a spectrum of complexity, yet both are considered species along the same Linnean continuum.

The existing toolset for describing pangenomes only tangentially measures this complexity^9,10^. In this paper, we measure complexity directly from the primary sequence itself rather than its myriad annotations. To do this, we transition to formal information theoretic definitions of complexity that operate on sequence ensembles^11,12^. The world is replete with sequence ensembles, linear strings of information that code sequential observations for numerous phenomena. Linguistic corpora^13^, neural spike trains^14^, and the microstates that feed statistical mechanics^15^ can all be described as sequence ensembles. Here, our sequence ensemble is the set of genomes used to describe a species. We propose complexity metrics that operate on these genomes directly, using unaligned and unannotated sequence. Our approach eliminates the need for an entire suite of bioinformatics tools that are costly to compute, over-parametrized, and often reference-bound. We show that our pangenome complexity definitions sort species into expected bacterial lifestyle categories previously assigned by more onerous pangenome based metrics. And we demonstrate that the term “species” must accommodate a startling range of pangenome complexity.

### Implementation

We measure complexity using the approaches highlighted in Figure 1. Our measures are all rooted in the concept of information entropy. Entropy measures the uncertainty of a probability distribution^16^. We measure entropy as

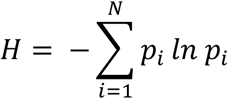

**Figure 1.**
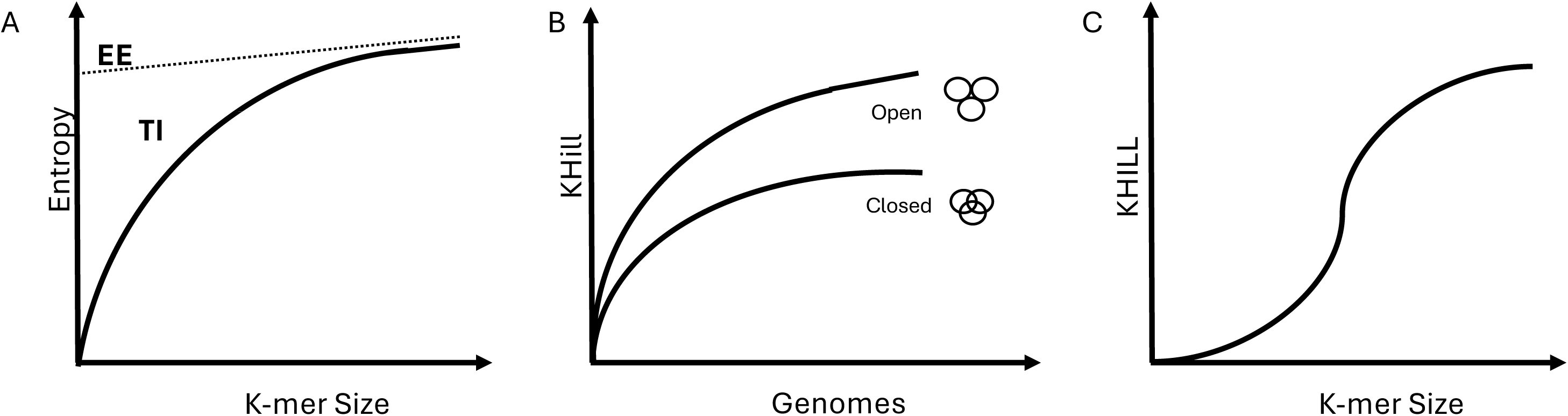
KHILL compression across all microbial taxa. We show KHILL values for all 10,510 RefSeq microbial taxa with at least two genome sequences as a semi-log plot. The red line plots y = x, the KHILL upper bound. The inset shows that the relationship is a power law distribution. Pathogens dominate the long tail, with tens of thousands of genomes compressing to a few genomic equivalents. High KHILL values are dominated by poorly characterized NCBI taxa and information rich species with complex pangenomes.

where *N* is the set of all possible states, *i,* and *p_i_* is the probability of the *i*th state. In fields where observations are organized as a set of linear strings, entropy can be calculated across blocks (subsequences) of various sizes. Plots of ensemble entropy against block size contain information about the ensemble’s complexity and compressability^11^ (Figure 1A).

These subsequences are called k-mers in the bioinformatics literature. While in complex ensembles, entropy always increases with k-mer size, this increase is often limited. A simple system will have more ensemble structure and lower entropy, whereas complexity implies less redundancy across block sizes and a higher entropy. Entropy growth curves measure how much surprise remains as you increase the k-mer size. An asymptote indicates full information capture. Continued growth at higher k-mer sizes implies there is more complexity to be learned.

In complex ensembles, we observe increased entropy because elements are unevenly distributed. For example, if a variable contains a highly predictable distribution, like a pangenome with a very tight genomic core, it’s entropy will be low and its information content high. Accessory genes increase surprise reducing predictable information. To measure these effects in a gene agnostic way, we analyze changes in information profiles along the block entropy growth curve^11^.

For every entropy curve, we calculate three quantities: the source entropy (*H*μ), excess entropy (EE), and transient information (TI) (Figure 1A). *H*μ is the ensemble randomness that remains after all the redundancies of longer and longer sequence blocks have been modelled. The y-intercept of the line chosen to approximate *H*μ is the excess entropy, or the intrinsic redundancy of the ensemble. The excess entropy is the non-random information capacity of our ensemble, information that can be captured because of its correlations. Finally, transient information is the amount of information that must be invested to discover *H*μ and fill out our excess entropy to its maximum. It is calculated as the area between the block entropy curve and the line describing *H*μ.

Genome ensembles with higher excess entropy and higher transient information are generally more complex. In our framework, this complexity is associated with a pangenome’s high fraction of accessory genes and a proportionately smaller core. Open pangenomes therefore have higher quantities of useful information (excess entropy) manifest as a larger set of accessory elements, but require more sequenced strains (transient information) to decode.

While entropy is key to understanding the information capacity of a system, our view of pangenome complexity requires an information theoretic analog to gene diversity. In previous work, we introduced KHILL, or the effective number of genomes, as a measure of complexity and structure in a viral pangenome^17^. The KHILL metric is an adaptation of beta diversity, an ecological concept used to measure species diversity across samples^18,19^. In ecological studies, researchers will often divide an area of interest into transects and count the number of species within each sample. The goal is to estimate overall diversity and to assess the efficacy of sampling. If the samples have many species in common at similar occurrence levels, the diversity difference between samples is low, and we calculate a lower effective number of samples or lower Hill number^20^. Hill numbers of this type provide an intuitive metric to compare species diversity across environments.

In a genomic context, we shift our frame of reference to measure information diversity. Rather than species, we count k-mers, and rather than transects, we sample genomes. We measure KHILL as

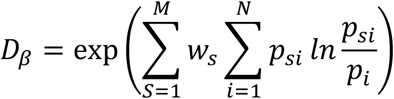

where *N* is the set of all k-mers, *M*, the set of all genomes, *p_si_* is the frequency of a particular k-mer *i* in a particular genome *s, p_i_* is the frequency of a particular k-mer *i* across all samples, and *w_s_* weighs all k-mers in sample *s* relative to all individuals counted in the experiment. Whereas in ecology, this quantity measures the effective number of samples, in genomics, the expression yields the effective number of genomes. KHILL summarizes the information diversity of any pangenome. A higher KHILL indicates a diverse set of k-mers, while a lower KHILL implies homogeneity. KHILL varies between 1 and the number of genomes sampled. A KHILL of one implies perfect compression, the information equivalent of sampling only a single genome. A maximal value signals no intersecting information, an ensemble of unique strings. KHILL can be applied to the pangenome of any species. As a measure of relative entropy, KHILL is in the family of information theoretic measures that employ the Kullback-Leibler Divergence^21^ to measure compression. We find that ensemble compression is a good proxy for pangenome complexity^17^.

But how can we use KHILL to measure pangenome complexity in practice? Like traditional pangenome analysis, we calculate saturation curves as we choose random genomes from the species ensemble. Instead of novel genes or pangenome fluidity, we plot KHILL as a function of genome accumulation (Figure 1B). As genomes accumulate, we accrue less unique information. If almost every k-mer in a new genome has been seen before, the curve plateaus, the mark of a closed pangenome. If new genomes continue to impart new information to the whole, the pangenome is open.

Importantly, not every k-mer size will yield meaningful results. In Figure 1C, we show that at low k-mer sizes, KHILL shows near perfect compression. That is, small k-mers are all found at approximately equal frequencies across all members of a species ensemble, collapsing the KHILL to 1. It is only as k-mer sizes increase that KHILL inflates to a second steady state. Information diversity accrues only at larger k-mer sizes. The optimal k-mer size for meaningful compression will vary depending on species but always corresponds to the elbow in the block entropy curve (Figure 1A).

The complexity-based approach we’ve outlined here sacrifices all knowledge of gene presence and absence. Rather than modular linear sequences with annotated functional elements, this view of the genome envisions each isolate as a container of information. We argue that by removing all levels of abstraction, unaligned and unannotated information can provide unbiased measures of pangenome complexity that inform species delimitation. Information theoretic complexity is as close to the data as we can get. Our approach suffers from none of the biases that can plague alignment, annotation and tree-based techniques. Abandoning traditional bioinformatics also furnishes speed. Here we calculate KHILLs and block entropies across all genomes of all microbial species in RefSeq.

## Results and Discussion

### The KHILL distribution of all bacterial species in RefSeq

We downloaded all available microbial genomes from RefSeq^22^ version 223 (73,079 taxa). Figure 2 shows the semi-log KHILL distribution of the 10,510 species represented by two or more genomes (283,097 genomes in total). Each point is an NCBI taxon. The red line is an upper bound: a species on this line has a KHILL equal to the number of genomes analyzed. Species sitting on this line have genomes with no information overlap. Most species have a KHILL below five. But some have KHILLs that approach their genome count. The 7095 genomes of Mycobacterium tuberculosis compress to a mere 1.8 effective genomes. The 79 genomes of *Prochlorococcus marinus* yield a KHILL of 21.9, occupying a much larger information space. This complexity gap is known from pangenome studies^7,8^, but the intuitive nature of the KHILL metric drives the point home. *P. marinus* is ten times more information diverse than *M. tuberculosis*. But both are considered taxonomic species.

**Figure 2.**
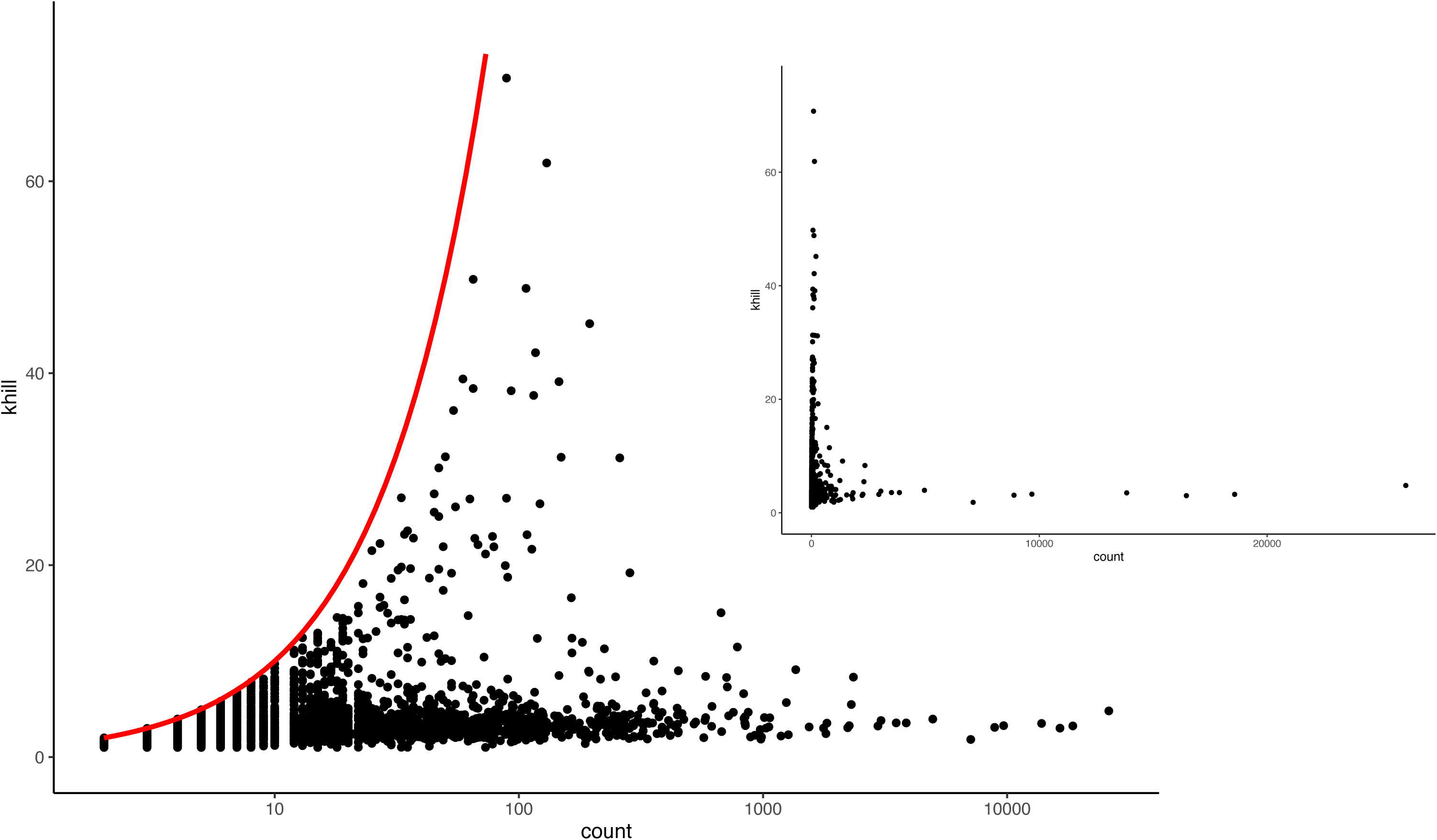
Pangenome accumulation curves. We show an alternative measure of the pangenome. Rather than counting the number of unique genes added with each genome in a set, we plot KHILL, a measure of information diversity. An open pan-genome’s KHILL will increase at a faster rate than a species with a closed pangenome (Panel A). The rate of increase in KHILL will vary depending on the evolutionary strategy and complexity of the pangenomes in question (Panel B). *H. pylori* is known to be recombinogenic, whereas *M tuberculosis* is clonal. In this framework all species shown are closed pangenomes after about 250 analyzed genomes except for the highly dynamic *H. pylori*.

We intentionally chose not to filter or screen any of this data. The plot is an unvarnished look at our current, accepted microbial species taxonomy. Clearly, the KHILL distribution of all microbial species follows a power law. The semi-logarithmic nature of our primary visualization allows us to separate some of the detail clumped at the origin. Most species have 10 or fewer genomes and seem to span their entire KHILL range. But most of the low genome count species hover between 1 and 2. Only 21 species have KHILLs higher than 20. These 21 species are represented on average by 82 genomes, a level of compression on par with *Prochlorococcus marinus*. The 28 species with more than one thousand genomes are more uniform and never eclipse a KHILL value of 10, perhaps reflecting the research attention they garner.

The extremes of this distribution reflect both a demonstration of and roadmap for research priorities. For example, the taxa with the three highest KHILLs are catch-all genus categories (*Flavobacterium sp*, *Pseudomonas sp*, and *Sphingomonas sp*). Poor taxonomic labels assigned to presumably challenging organisms are bound to combine divergent genomes. A genus grab-bag likely obscures reticulated species that require further sampling and genomic work. In contrast, species represented by thousands of genomes are among those that make humans most sick. *E. coli* is represented by 26,088 genomes, *K. pneumoniae* by 18,575. But their KHILLs are 4.81 and 3.24, respectively. The more than 25 thousand genomes of *E. coli* compress into just 5 effective units.

While this rate of genome accrual is likely the result of public health initiatives and funding priorities^23^, we show that representatives for these species appear to be highly redundant. That is, information is not accumulating at the same rate as genomes in the database. This is not meant to impugn pathogen genome sequencing projects, many of which are looking for subtle genomic signatures in strains or variants causing outbreaks. But low KHILLs among these pathogens imply that their pangenomes have saturated. In terms of their information structure, additional genome sequencing is not shedding any further light on the complexity of the species.

We view either end of this power law distribution as revealing key aspects of the culture of science: relentless classification of even the poorly understood, and relentless investment in the small group of organisms with an outsized impact on human welfare.

Despite these anthropocentric biases, the distribution in Figure 2 contains evidence that KHILL does highlight species with complex pangenomes. *Pseudomonas fluorescens* (KHILL = 45.2), a versatile, free-living, gram-negative bacteria found in soils and water is known to have a large, complex pangenome^24^. And *Buchnera aphidicola* (KHILL = 48.8), an obligate endosymbiont of pea aphids is known to have an unusual amount of genome divergence^25^.

Thus far, we have compressed the signal from all available genomes for each species into a single point. In the next section, we instead calculate KHILL saturation dynamics. Accumulation curves have long been used in ecology to measure diversity in species distributions^26–28^. These curves can also quantify string diversity in sequence libraries^29^. In the next section, we apply this concept to pangenomes.

### KHILL accumulation curves

In their seminal work in *Streptococcus agalactiae*, Tettelin et al^4^ showed that every new sequenced strain contained new genes. This became the hallmark of an open pangenome. Though their accumulation curve exhibited saturating behavior for five or fewer genomes, thereafter the curve rose linearly with no clear asymptote. Genes seemed unable to demarcate a clear taxonomic boundary.

Still, genes are intuitive functional units for pangenome analysis. Most programs available for pangenomes are built around a gene-centric view of biological information^9,10^. But a reliance on genes requires faith in their annotation, tolerates bias towards model organisms, and relegates non-coding elements. Computing on genes is also costly^30^. And because assigning gene orthology is difficult, most existing pangenome tools require hundreds of CPU hours to analyze thousands of genomes. As an alternative, in Figure 3A, we show KHILL accumulation curves.

**Figure 3.**
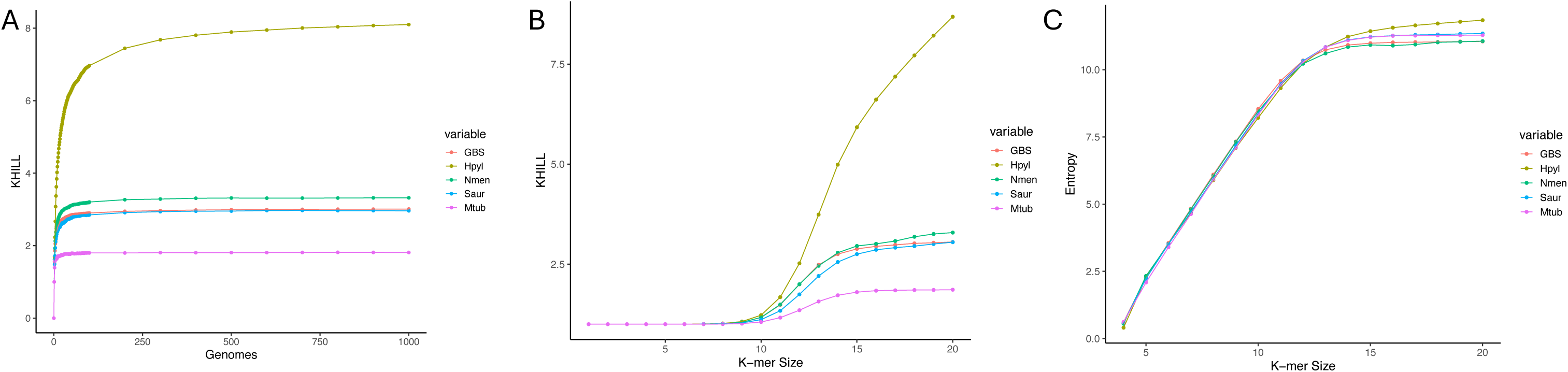
Varying k-mer size. Increasing k-mer size increases both KHILL and entropy. KHILL increases rapidly at the k-mer size that corresponds to the entropy elbow (A). The block entropy curve (B) yields two important insights into the information content of an ensemble. The y-intercept of the approximate asymptote is the excess entropy, a quantity often interpreted as the computational capacity of the system. The area between the line tracing this asymptote and the curve itself is the transient information, the information cost required to synchronize to the steady-state entropy of the ensemble. For our five pathogens, we show that all but H. pylori settle into a KHILL maximum (C), and that even small changes in the block entropy curve (D) can lead to large differences in information diversity as reflected by KHILL.

KHILL decenters the gene and instead focuses on sequence information, distilling signal into genome equivalents. KHILL accumulation curves build out a sequence ensemble. Our sequence ensembles are the set of all genomes that enrich our picture of any given species. Each genome increases the ensemble’s complexity, but at differing rates depending on the organism. Like gene-based approaches, if a pangenome is closed, KHILL will eventually flatline. But if new genomes consistently deliver novel information, KHILL will never asymptote. KHILL operates as an information-based, computationally tractable proxy of more cumbersome metrics like pangenome fluidity^5^. The two quantities display a linear relationship (Supplemental Figure 1) but are orders of magnitude apart in terms of computational burden (Supplemental Figure 2).

Genome size is known to be a confounding factor in orthology-based pangenome analyses^6,31,32^. In Supplemental Figure 3A, we show that the relationship between genome size and pangenome fluidity is linear. The influence of genome size on KHILL, however, is less pronounced (Supplemental Figure 3B). We built linear models of this relationship, separating fitted from residual components. KHILL fitted to genome size flatlines in plots against pangenome fluidity, implying that genome size is not driving the relationship between the two quantities (Supplemental Figure 4A). The KHILL residuals – the component unmodeled by size – correlate with pangenome fluidity (Supplemental Figure 4B).

In Figure 3A, we show KHILL accumulation curves for five pathogenic species. Unlike conclusions drawn in the Tettelin^4^ paper, the *Streptococcus agalactiae* pangenome seems to be closed with an asymptote at about 3 effective genomes. In keeping with its reputation as a clonal organism with no evidence of recombination^33^, *Mycobacterium tuberculosis* levels at a KHILL of 1.8. *Staphylococcus aureus* and *Neisseria meningitis* asymptote at KHILLs of 2.4 and 2.6, respectively. *S. agalactiae*, *M. tuberculosis, S. aureus,* and *N. meningitis* all seem to require only about 150 randomly selected GenBank strains to confirm a closed pangenome. Among the five, only *Helicobacter pylori*, an organism known to exhibit intense recombination^34^, is an open pangenome. Even the thousandth *H. pylori* genome adds significant unique information. *H. pylori’s* high KHILL suggests a complex pangenome. Differential rates of saturation to various KHILL maxima across these species suggests a new kind of comparative genomics.

### KHILL across k-mer sizes

KHILL requires that we set a k-mer size. Shorter k-mers enhance compression but sacrifice combinatorial nuance^17^. Longer k-mers capture more interaction information but grow exponentially in computational difficulty. At the maximum theoretical extreme, as k-mers approach the size of the genome, KHILL balloons to the number of input sequences. Entropy also increases with k-mer size^11^. Given a sequence ensemble, calculating entropies and KHILLs with increasing k-mer size spawns new measures for species information content and prescribes a string size required to make that information content bloom.

KHILLs at small k-mer sizes collapse all available information in the ensemble. But as we increase block size, KHILL moves through a steep transition to a second steady state. K-mer sizes beyond this transition increase KHILL at a slower rate. In Figure 3B, we show these curves for our five pathogens. All organisms except *H. pylori* exhibit dual steady-state dynamics. Though the rate of increase appears to abate, *H. pylori* KHILL continues to increase even beyond a k-mer size of 21.

If an ensemble contains information restricted to only subsets of strains – the hallmark of an open pangenome – we would expect KHILL to continue its increase over larger k-mer sizes. In the case of non-recombining organisms with closed pangenomes, like *M. tuberculosis*, the k-mer ensemble stabilizes across all available strains by a k-mer size of 15. Since open pangenomes maintain diverse populations of information restricted to subclades, KHILLs of these organisms will likely never converge regardless of string size.

### Block entropy curves

Block entropy curves measure entropy as a function of k-mer size. Larger k-mers generate larger entropies. But in genome ensembles, entropy generally settles into a steady-state after a k-mer size of about 15 (Figure 3B). K-mer sizes at the block entropy elbow echo those needed to achieve a steady KHILL. In other words, the rate of entropy increase abates just as KHILL transitions between its two steady states (Figure 3C). Diminishing returns with higher string sizes suggests a k-mer optimum. Given an optimal string size, we show that entropy captures the information capacity of our ensemble and KHILL measures its information diversity. The two metrics together join to quantify its complexity.

Block entropy curves are key to complexity analysis in other fields. In physics, sequences of molecular states can describe phase transitions in materials at different temperatures^15^. In neuroscience, sequences of action potentials can measure collective behavior in neural networks^35^. We borrow these concepts for the analysis of genome sequences. After a linear ascent common to most sets of genomes, entropy plateaus to the system’s source rate, *Hμ*.*Hμ* measures the intrinsic randomness in the sequence ensemble, the noise that achieves no structure regardless of k-mer size^11^. In practice, *Hμ* is only defined as k-mers approach the size of the genome. For more reasonable and calculable k-mer sizes, we approximate *Hμ* by measuring the slope of the entropy curve at the largest k-mer sizes we can manage.

The y-intercept of this line is an estimate of the excess entropy (EE)^11^. The excess entropy has been called the predictive information^12^, the source’s memory, or its redundancy. Excess entropy is the system’s useful complex structure. *Hμ* is its inevitable noise. Here, we interpret excess entropy as the computational capacity of the system, the capacity of the pangenome to code useful functions for a given environment. As block entropy curves reach a steady state, we approach true values for *Hμ* and excess entropy. Though much more subtle than KHILL, we show a higher *Hμ* and higher excess entropy in *H. pylori* than the four other pathogens shown in Figure 3C. A higher excess entropy implies higher pangenome complexity.

Different ensembles will saturate at different rates and at different k-mer sizes. The process of synchronizing to the steady-state is also a measure of complexity, and one we can approximate without exploring k-mer sizes in the extreme. The transient information (TI)^11^ is the amount of information we must learn as we discover the structure of an ensemble (Figure 3C). TI measures our information investment as we probe the dynamics of pangenome evolution.

Species with large TIs require more information to discern their internal structure. Like pangenome fluidity, TI trends upwards with genome size (Supplemental Figure 5). A linear model of TI with respect to genome size establishes that the residuals track more closely with pangenome fluidity (Supplemental Figure 6). Genome size is therefore factorable in our view of pangenome evolution. With TI, we show that learning complexity is its own measure of complexity.

Recombination and mutation are two primary drivers of microbial evolution. In Figure 4, we simulate 100 1 Mb genomes for 25 combinations of homologous recombination and mutation rate. Both EE and TI increase as mutation rate and recombination rate increase. In other words, the information capacity and the amount of information required to measure that capacity increase with both modes of sequence change. As we have previously demonstrated elsewhere^30^, recombination seems to exert the larger effect.

**Figure 4.**
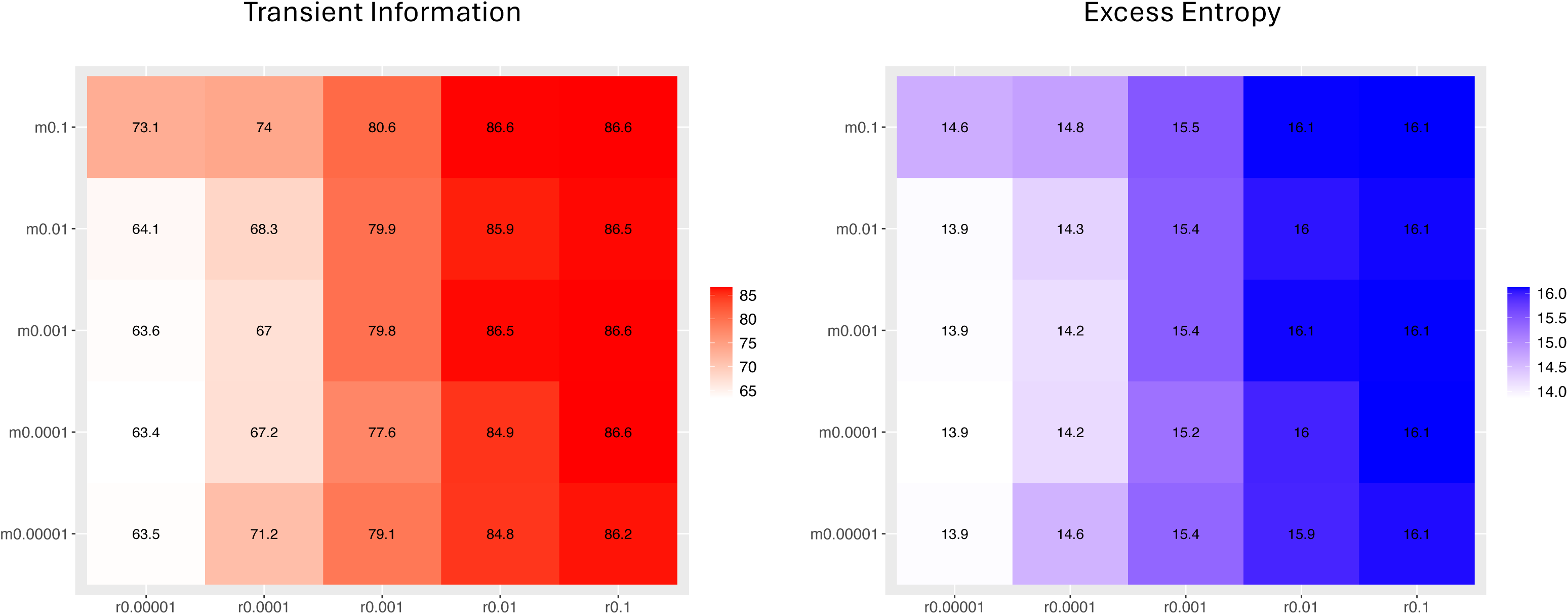
Simulations. We simulated 10 genomes for every recombination rate and mutation rate pair. Higher rates of either form of genomic change increase both the transient information and the excess entropy, two hallmarks of increased ensemble complexity.

### Block entropy and KHILL track bacterial lifestyle

The pathogens we analyze in Figure 3 are all bacteria subject to one environment: the interstitial spaces of the human body. What if we instead look at free living bacteria? Or obligate intracellular microbes? In Figure 5, we show block entropy (A) and KHILL (B) curves for 10 random Genbank genomes of *Pseudomonas fluorescens*, *Rickettsia japonica* and *Staphylococcus aureus*. *P fluorescens* is found in profoundly variable environments^24^. *R. japonica* is non-motile and restricted to living in the cytoplasm of its eukaryotic host^36^. In terms of both block entropy and KHILL, we see a gulf in complexity between these two organisms.

**Figure 5.**
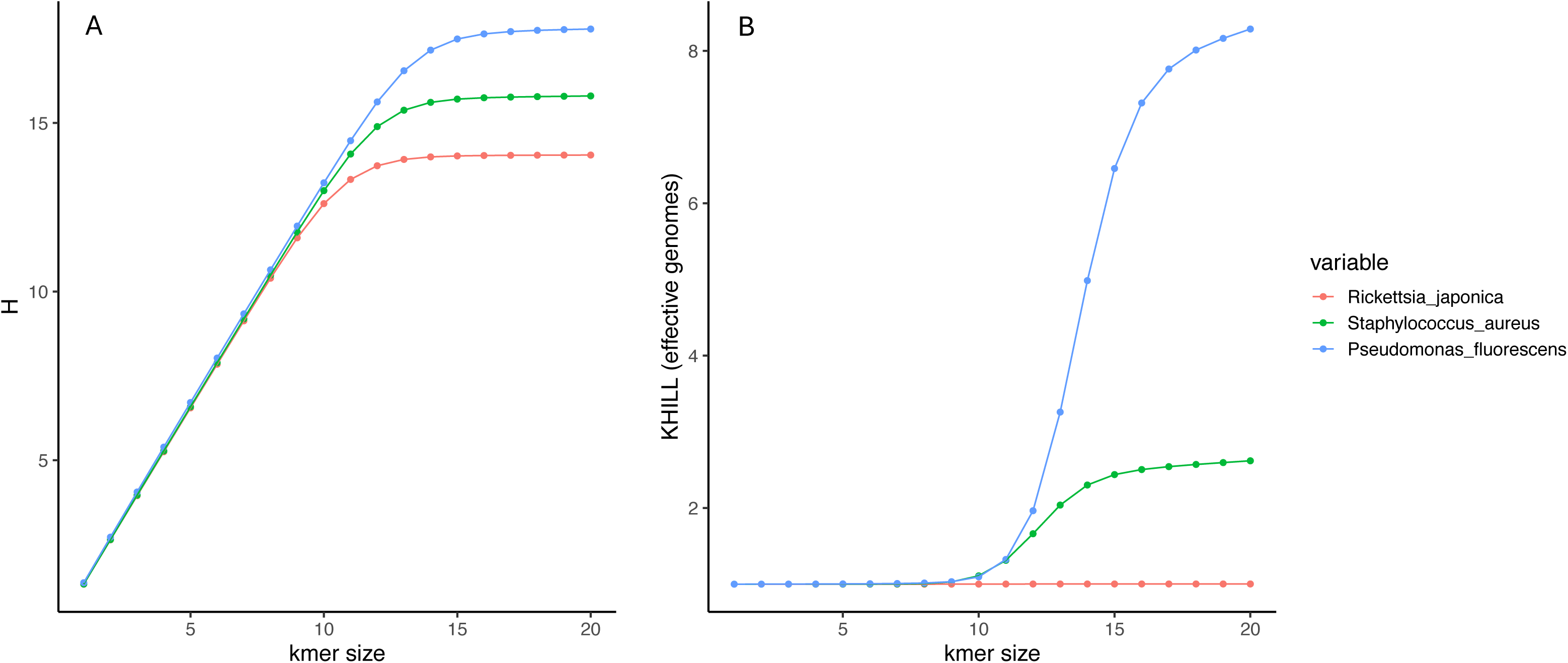
Lifestyle extremes. We show block entropy curves (A) and KHILL curves (B) with increasing k-mer sizes across three organisms with very different lifestyles. *P fluorescens* is a cosmopolitan, free living microbe. *R. japonica* is an endosymbiont restricted to the cytoplasm of a single host. These lifestyle differences are reflected in their information capacity (A) and information diversity (B) extremes. *S. aureus* a largely clonal pathogen is an example of an organism that relies on a host for propagation but can survive in the environment for short periods of time.

In the complexity literature, the first and second derivatives of the block entropy curve provide information about the excess entropy and the k-mer size that achieves peak predictability, respectively^11^. If excess entropy is the ensemble’s store of useful information, peak predictability is the string size most useful to discern that information. For our application, we interpret this as the minimum k-mer size required to retrieve information about the system. The curves in Figure 5 suggest that species with grossly different lifestyles may require very different k-mer sizes to optimize information about their pangenome complexity (Supplemental Figure 7). These curves also hint at a new method to select k-mer sizes that maximize predictability, perhaps clarifying size selection procedures in short read alignment and assembly^37,38^. *P fluorescens* requires a k-mer size of 14 to coax peak information out of the ensemble. For *R. japonica* a k-mer size of 12 suffices.

The complexity gap between these two extreme organisms is in keeping with known differences in pangenome diversity and reflects the stark contrast in the environmental challenges they face. We selected *S. aureus* as intermediate between these two lifestyle poles. Remarkably, at a k-mer size of 20, 10 strains of *P fluorescens* contain the information equivalent of over 8 genomes, while all 10 R*. japonica* collapse to a KHILL of 1 across the entire k-mer expanse.

Between the flexible cosmopolitan (*P fluorescens)* and the cytoplasmic shut-in (R*. japonica*), bacteria have settled into several intermediate lifestyle traits. Some are obligate, some facultative. Some move and some do not. Some are pathogens and others, mutualists. Dewar et al^6^ recently annotated 126 bacterial species and found that more fluid pangenomes are usually free living, facultative, motile, extracellular, and non-pathogenic. Variable environments seem to require genomic flexibility. On the other hand, obligate, host-bound pathogens tend to have a streamlined pangenome.

Figure 6 shows KHILL distributions of Dewar et al’s 126 species with respect to their pathogenicity. We show that pathogens have a significantly lower KHILL than mutualists, a finding that recapitulates results seen with more expensive metrics like pangenome fluidity. In Figure 7, we show that genome size corrected residuals of transient information also correspond to our intuition regarding pathogenic organisms. Pathogens face homogenous environments. Their pangenomes are often less complex as a result.

**Figure 6.**
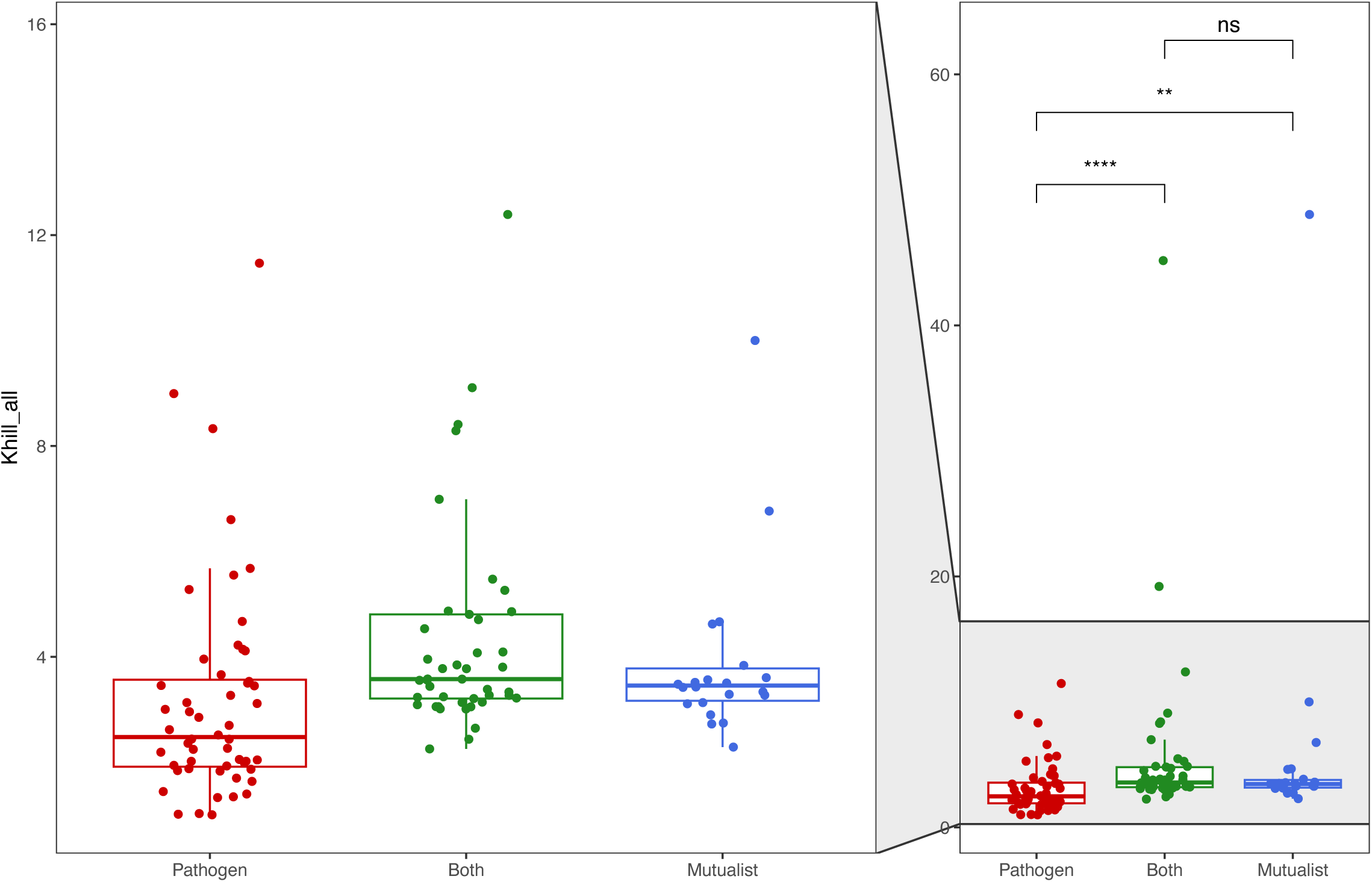
Pathogens have lower KHILLs. We show that the pathogens among the 126 species curated by Dewar et al have significantly lower KHILLs than mutualists or organisms that both infect their hosts and live independently. Pathogens exist in the controlled space of their hosts, subject to few disturbances. Their low information diversity reflects a lack of accessory genes that might be more functional among organisms that must contend with dynamic environments.

**Figure 7.**
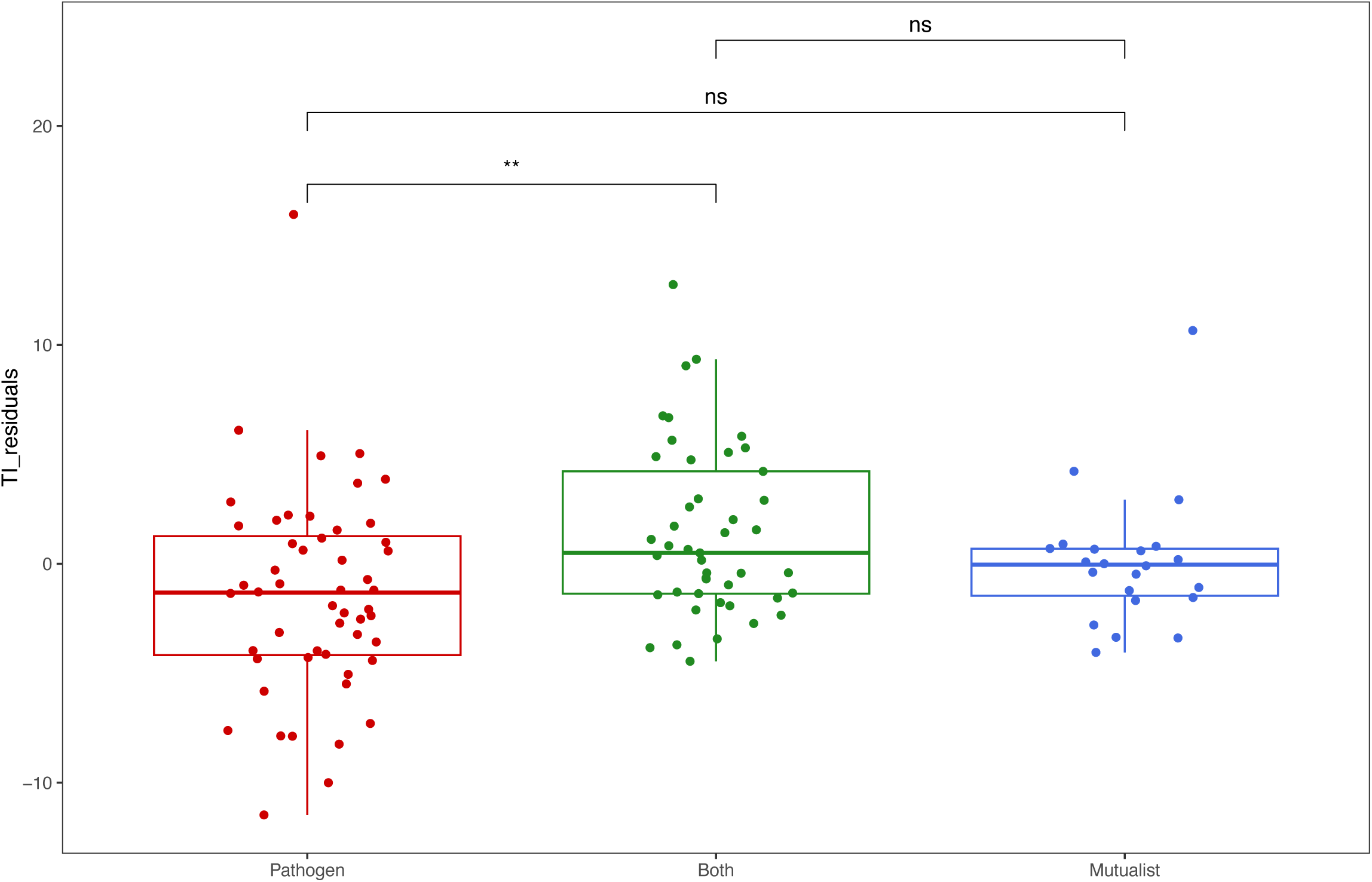
Pathogens have lower transient information. We show that pathogens have lower transient information implying a smaller information capacity relative to non-pathogenic organisms. Here we show transient information residuals after a genome size correction.

KHILL distributions sort with our expectations on every other lifestyle trait annotated by Dewar et al. Bacteria that are host-bound, intracellular, obligate, and/or non-motile all have significantly lower KHILL than their free-living, extracellular, facultative, and motile counterparts (Supplemental Figure 8). Using linear models of both KHILL and TI against genome size, we confirm that it is the residuals of both metrics that undergird our measurement of pangenome complexity (Supplemental Figures 9 and 10). Residuals of transient information agree with our expectations regarding lifestyle but the distinctions between traits are less resolved (Supplemental Figure 11). This is in keeping with the higher resolution of KHILL (Figures 3 and 5). For both motility and host reliance, TI residuals show no distinction between populations.

### Conclusion

The information-based techniques we introduce here are lightweight and capable of handling orders of magnitude more data than traditional pan-genomic approaches. We do this without sacrificing the biological signal we expect. Our metrics track with bacterial lifestyle. Microbes enmeshed in variable environments have higher KHILLs and higher transient information. Both are markers of pangenome complexity, and both are derived from a formal literature that quantifies complex systems. We find that compressibility is a viable comparative strategy. KHILL combines the effects of genetic distance and population heterogeneity.

Transient Information is a useful proxy for species information capacity. We use both metrics here to explore bacterial lifestyle, standardize molecular species definitions, highlight tenuous classifications, and confirm pangenome saturation.

## Methods

### The program

We calculate *Hμ*, excess entropy, and transient information from block entropy curves using the program khill_complexity available at https://github.com/narechan/khill_complexity. All sequences in an ensemble are provided as independent fasta files in a single input directory. The program will accept either a single k-mer size or the range of k-mer sizes required of block entropy curves. For every k-mer size, the program also generates a KHILL value that quantifies the level of genome compression.

We provide two mechanisms to increase the speed of these calculations. First, reading FASTA files and calculating KHILL values is parallelized across all input genomes. The user can set how many processors they wish to dedicate to the calculation. Ideally, one per input genome sequence. Second, we provide a bottom-k hashing heuristic which allows us to sketch^39^ the input k-mers. For example, a rate of 10% means that the lowest tenth of the hash values (corresponding to k-mers hashed with murmurhash3) are sketched into the set of strings used for the block entropy curve and the KHILL calculations.

A typical khill_complexity run of 10 genomes would be invoked like this:

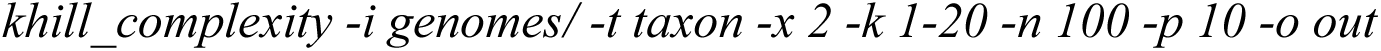

Here, the genomes (-i genomes) with their taxonomic designation (-t taxon) are run at k-mer sizes ranging 1 to 20 (-k 1-20) across 10 processors (-p 10). The program can be run with hashing (-x 2) or with straight k-mers (-x 1). If hashing is employed, the k-mers can be sketched (-n 100). At this setting, only the lowest 1% of hashes are included in the analysis.

### Refseq and clade analyses

We analyzed all bacterial taxa in RefSeq (version 223) with two or more genome representatives. We randomly selected up to 1000 genome assemblies from *Streptococcus agalactiae, Mycobacterium tuberculosis, Staphylococcus aureus, Neisseria meningitis,* and *Helicobacter pylori* for our analysis of pathogenic clades. We downloaded all genomes available in RefSeq for the 129 taxa selected in Dewar^6^ et al. We plotted KHILL and transient information across all their gross phenotypic categories: 1. pathogen/mutualist; 2. host-bound/free; 3. non-motile/motile; 4. intracellular/etracellular; and 5. obligate/facultative. We performed linear regressions in R across all the following pairs, modeling both fitted and residual components: 1. KHILL vs pangenome fluidity; 2. pangenome fluidity vs genome size; 3. KHILL vs genome size; and 4. transient information vs genome size.

### Simulations and annotations

We simulated 1000 1 Mbase genomes with varying mutation and homologous recombination rates in SimBac. For a recombination rate of 0.0001 and a mutation rate of 0.01we issued the following:

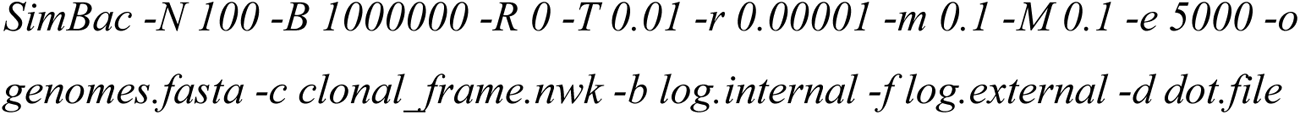

To compare our runtimes with traditional gene-centric pangenome methods we annotated and built orthologs for 100 random genomes from the 129 taxa found in Dewar et al. We used prokka^40^ to annotate and roary^10^ to build gene orthologs.

---

**Supplemental Figure S1.**
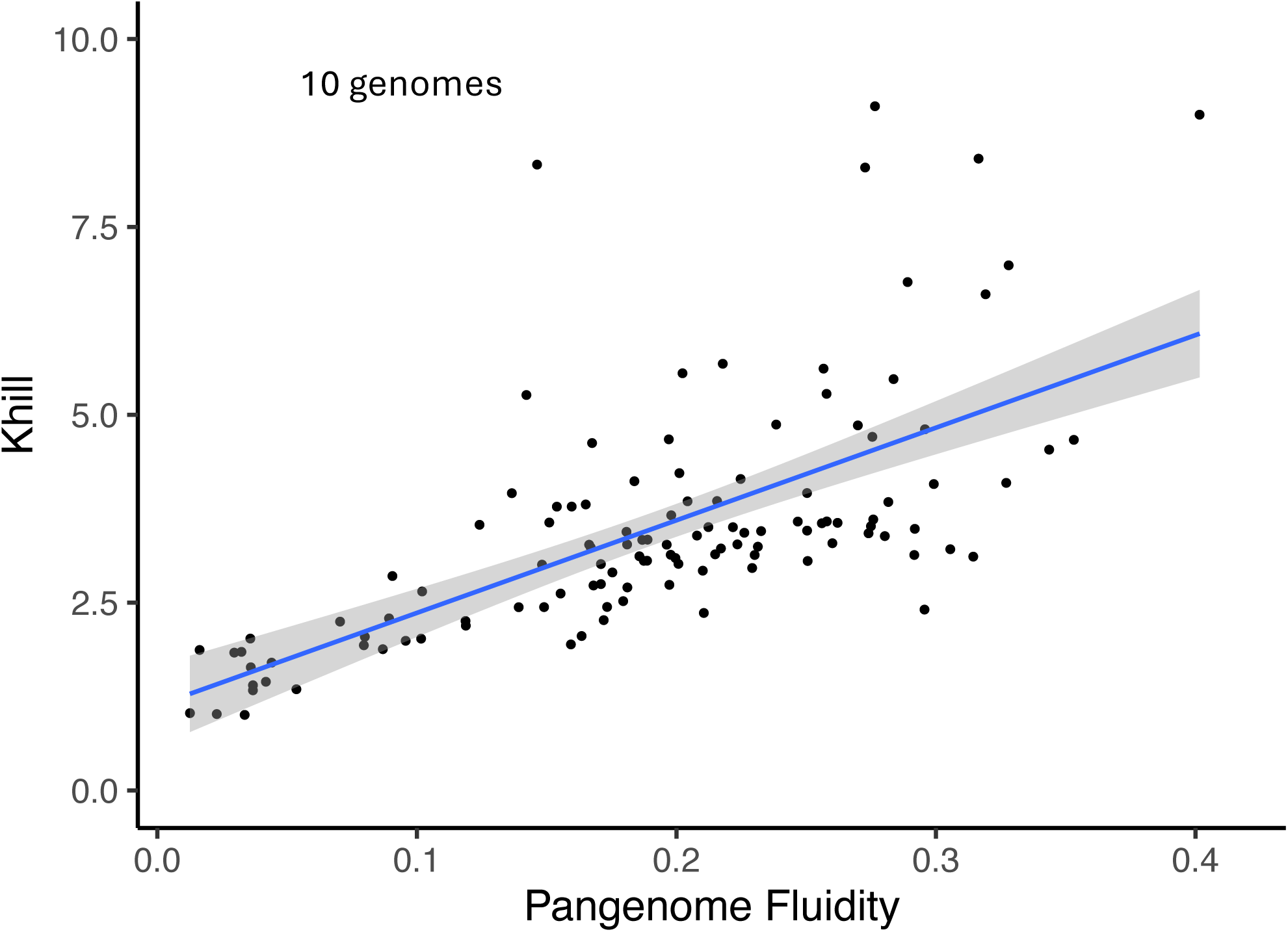
KHILL correlates with pangenome fluidity. We show that KHILL seems to have a linear relationship with pangenome fluidity.

**Supplemental Figure S2.**
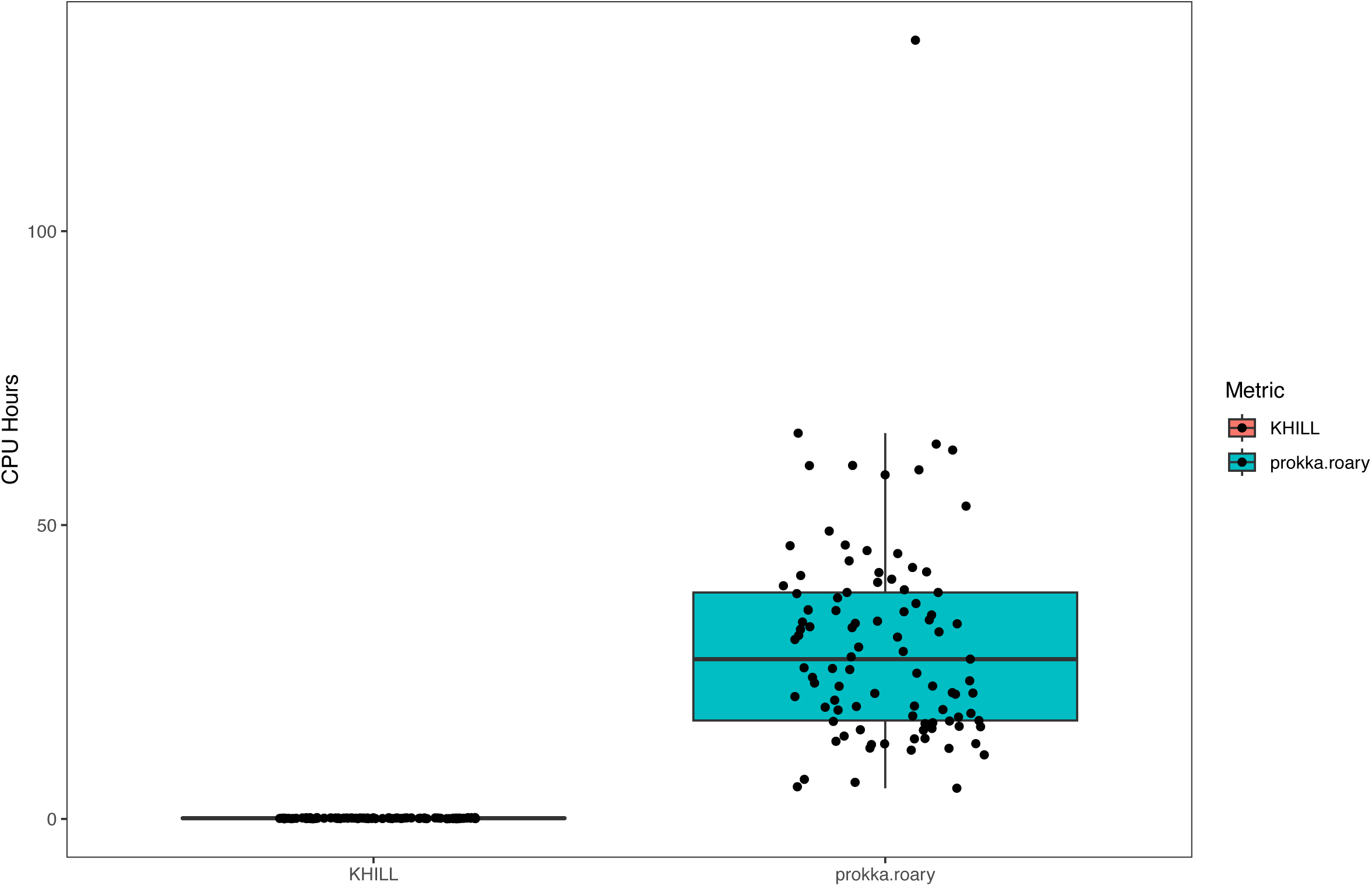
Benchmarking. We show that calculating KHILLs is orders of magnitude faster than traditional microbial pangenomic methods which require annotation (here by prokka^40^) and gene orthology (here calculated in roary^10^).

**Supplemental Figure S3.**
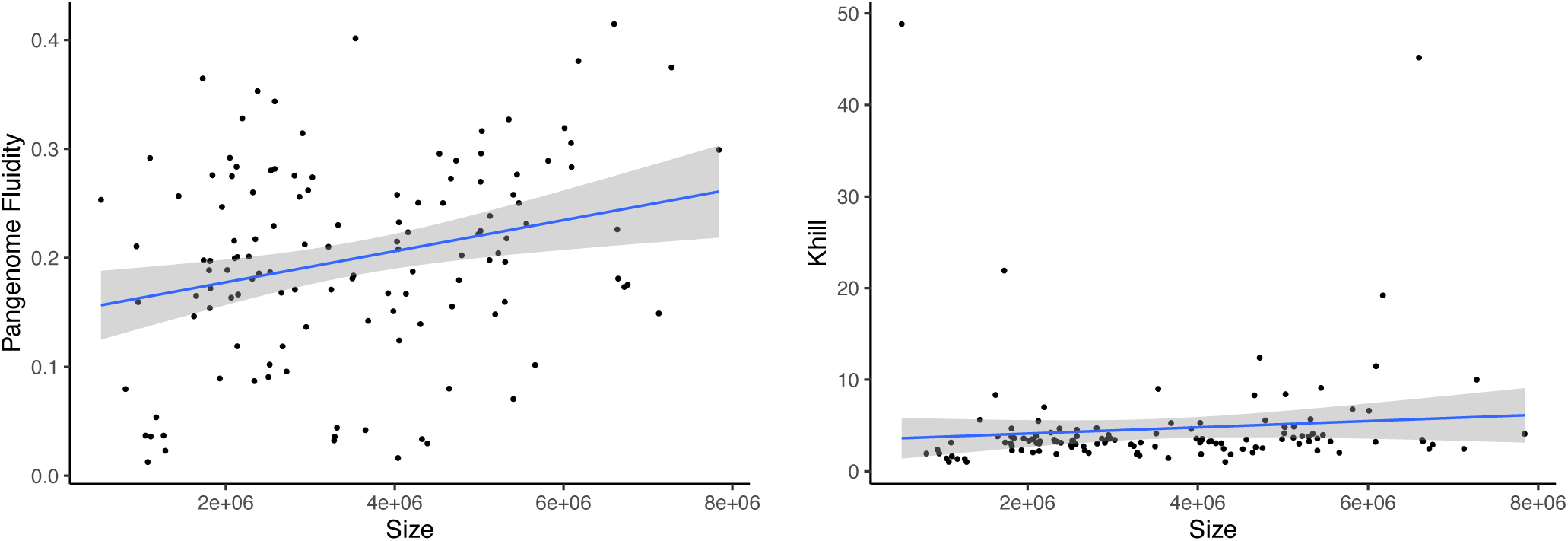
Genome size and KHILL. We show the relative effect of genome size on pangenome fluidity and KHILL.

**Supplemental Figure S4.**
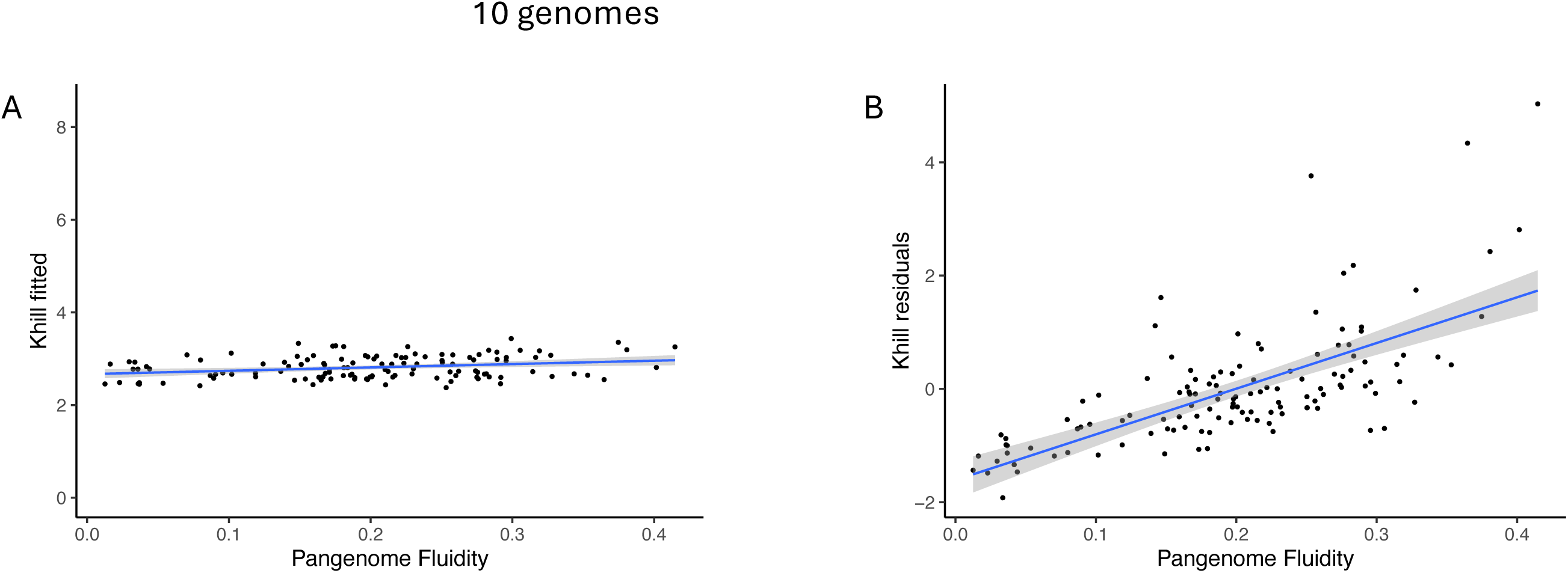
KHILL components. We calculate a linear model of KHILL with respect to genome size and calculate the fitted and residual components. The KHILL residuals drive KHILL’s correlation with pangenome fluidity suggesting that KHILL values are largely independent of genome size.

**Supplemental Figure S5.**
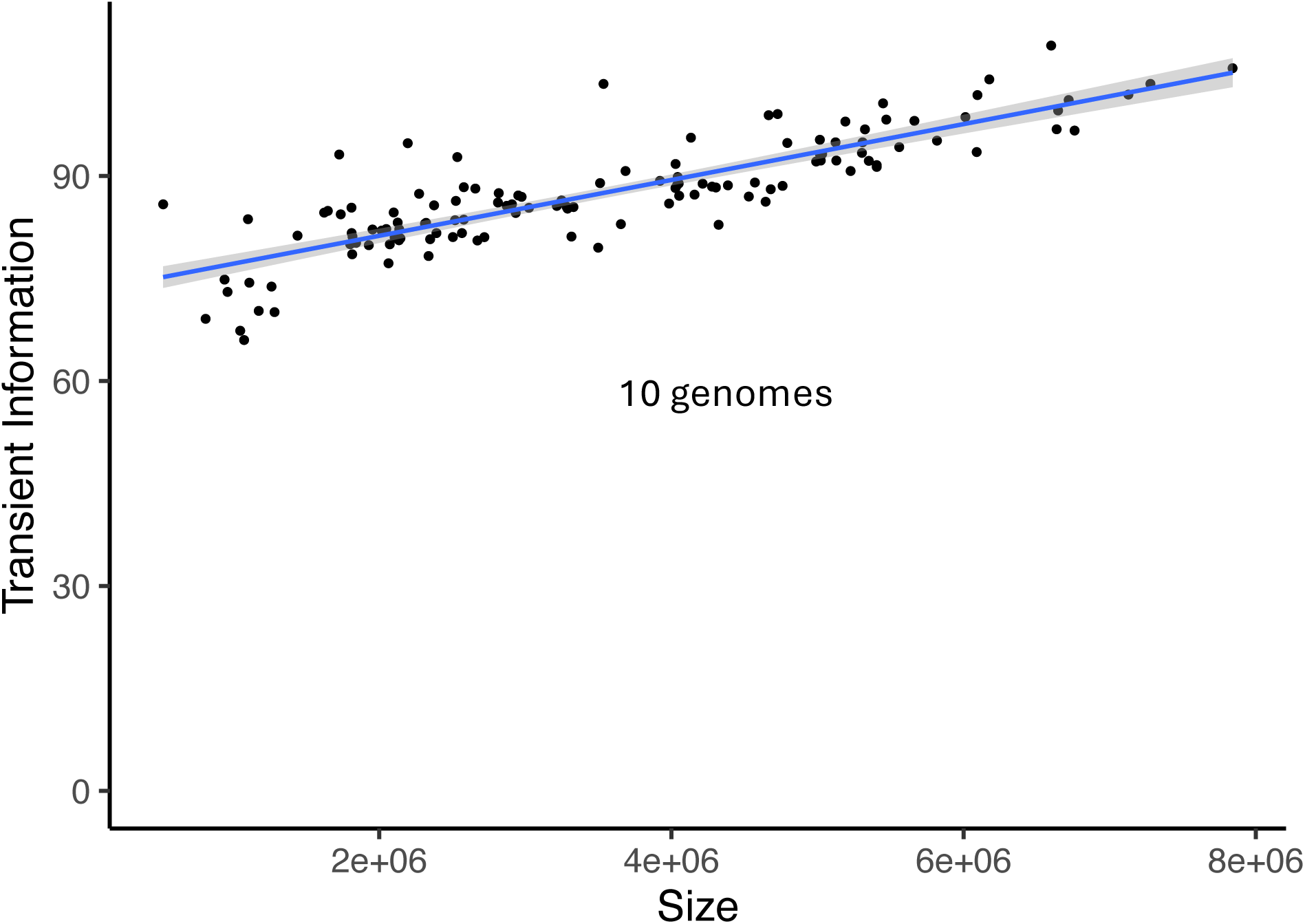
Genome size and transient information. We show that larger genome sizes are correlated with larger TI values.

**Supplemental Figure S6.**
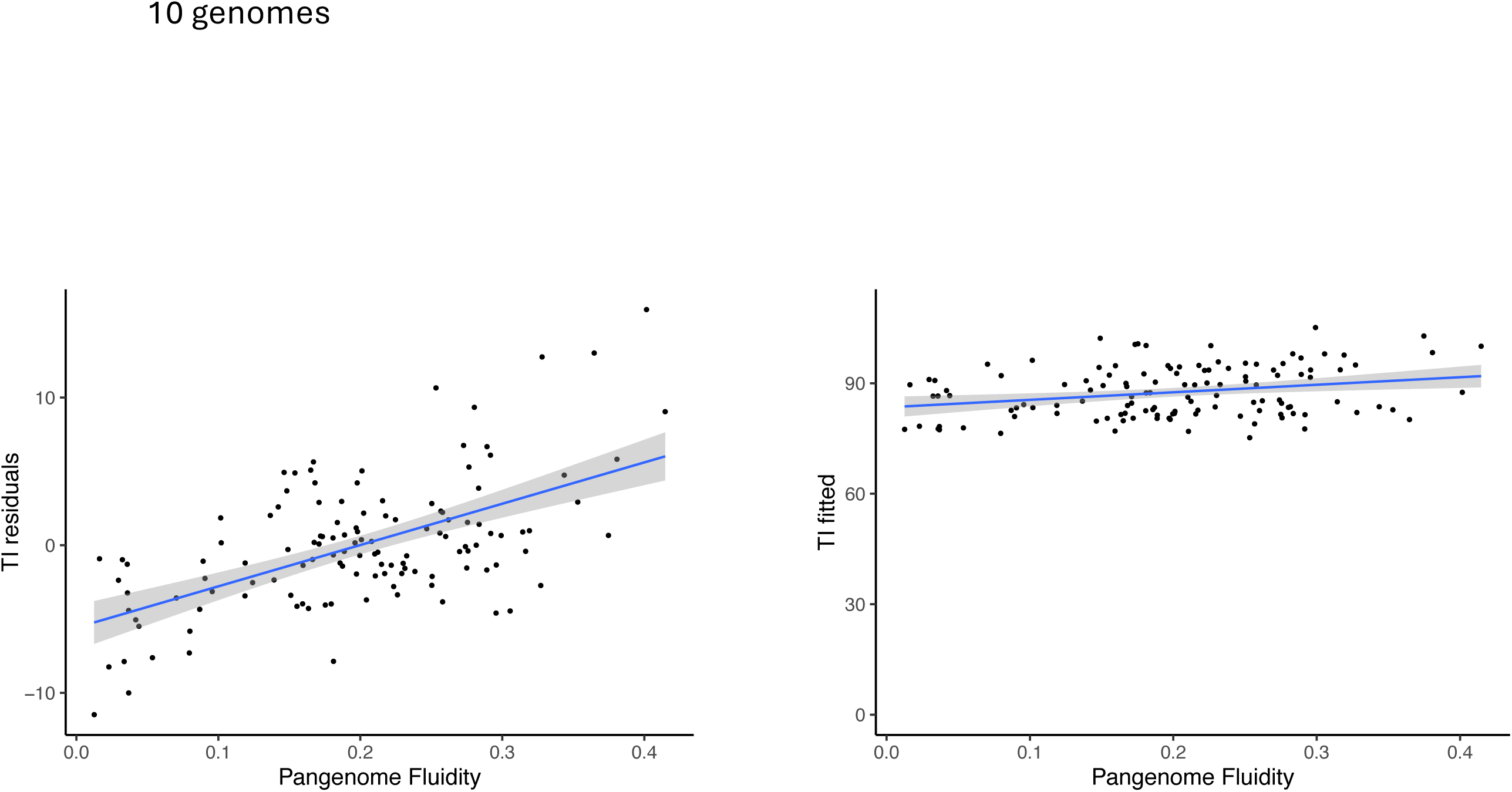
Transient information components. We calculate a linear model of Transient Information with respect to genome size. The residual components again largely drive correlation with pangenome fluidity.

**Supplemental Figure S7.**
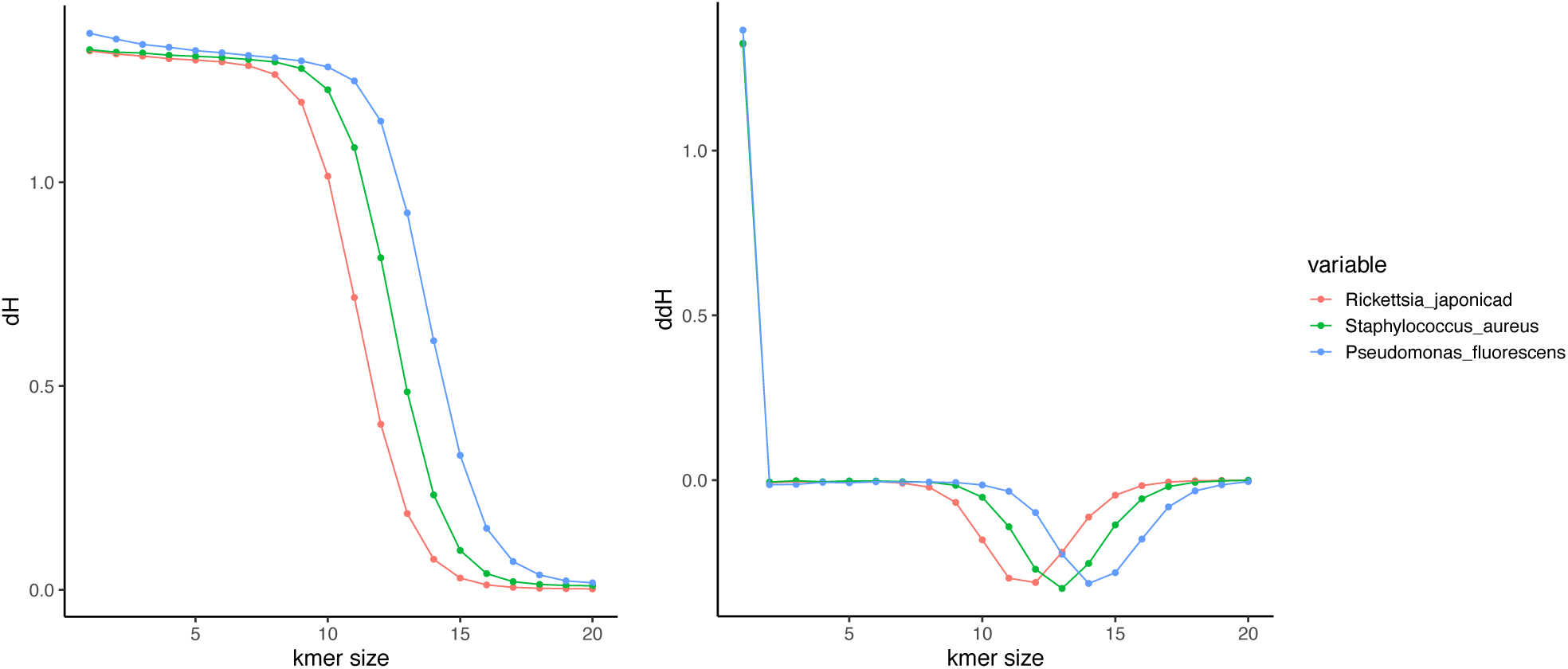
The first and second derivatives. We show the first and second derivatives of block entropy curves across three microbes with very different lifestyle traits. The first derivative is a useful and alternative way to model excess entropy. The minima on second derivative curves provide insight into the k-mer size that affords peak predictability for each sequence ensemble.

**Supplemental Figure S8.**
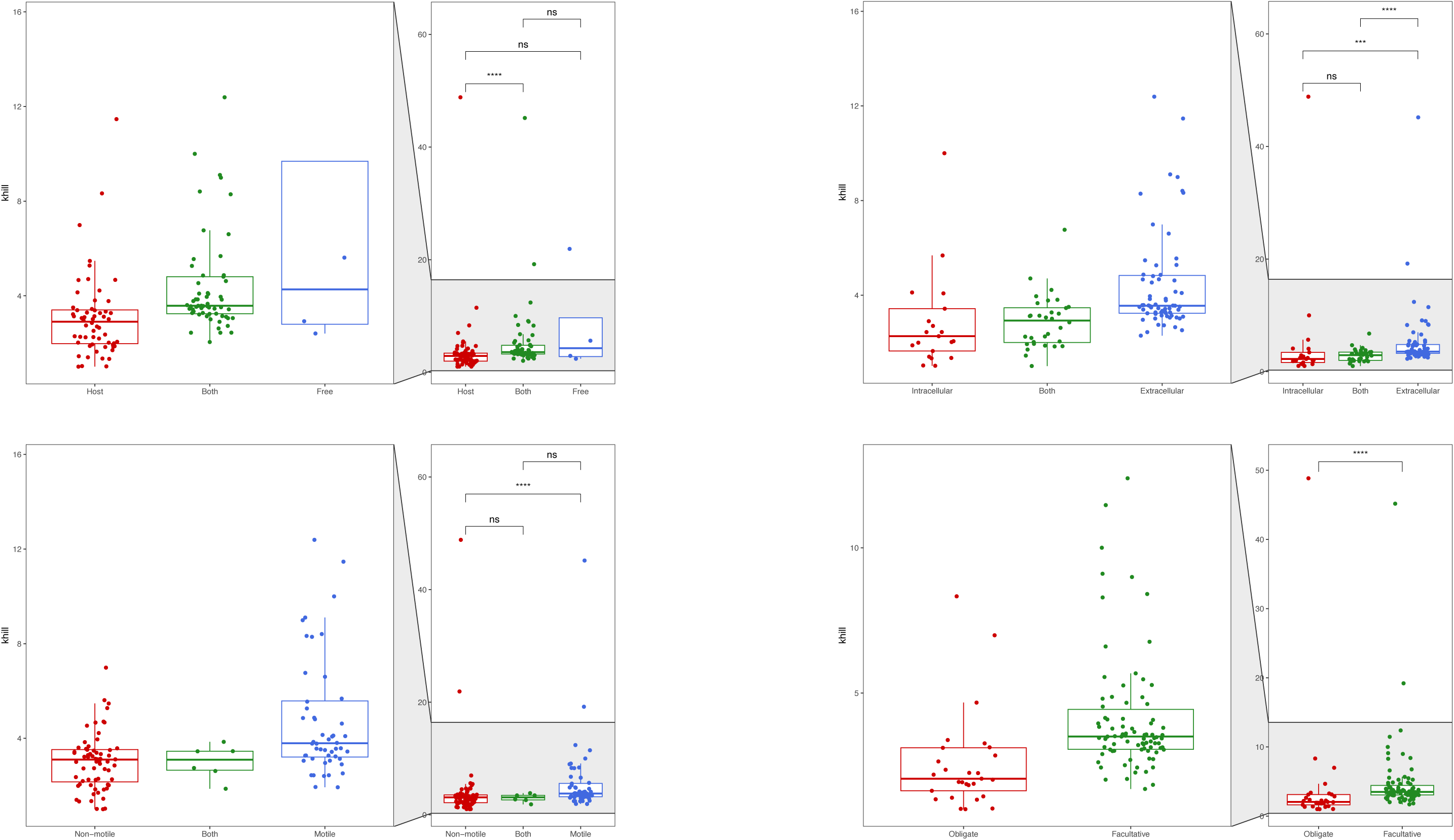
KHILL distributions against four lifestyle traits. We show significant KHILL differences between organisms that are host-bound and free living, intracellular versus extracellular, non-motile versus motile, and obligate versus facultative.

**Supplemental Figure S9.**
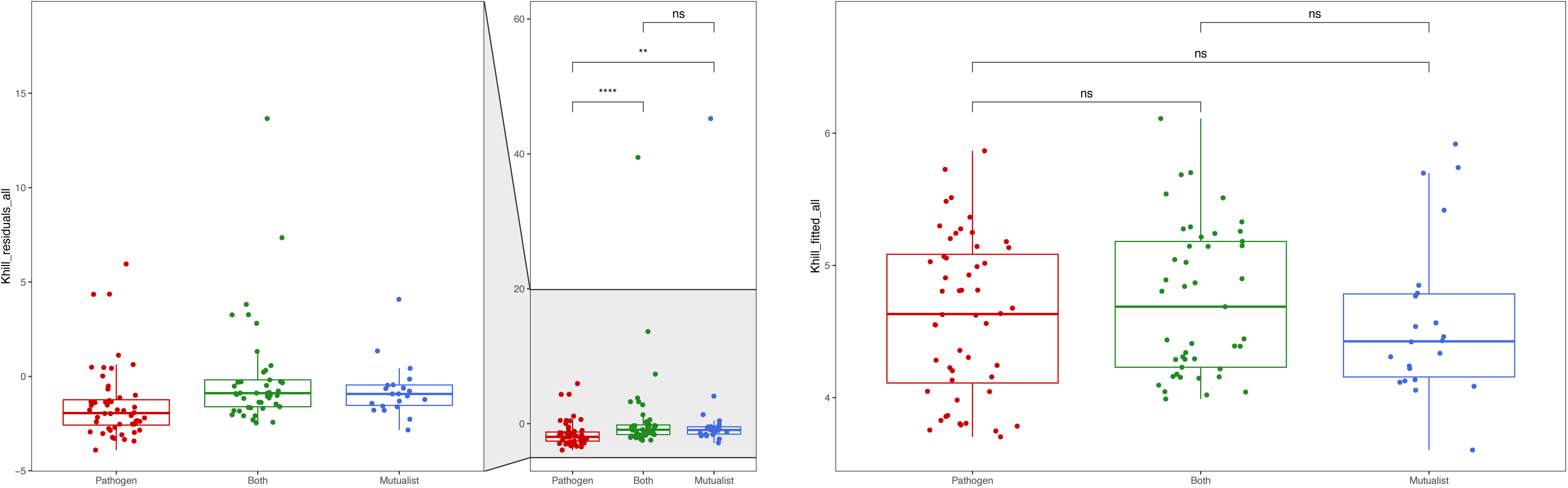
KHILL components against pathogenicity. We show that KHILL residuals from a linear model against genome size separate pathogens from mutualists. The fitted components yield non-significant results.

**Supplemental Figure S10.**
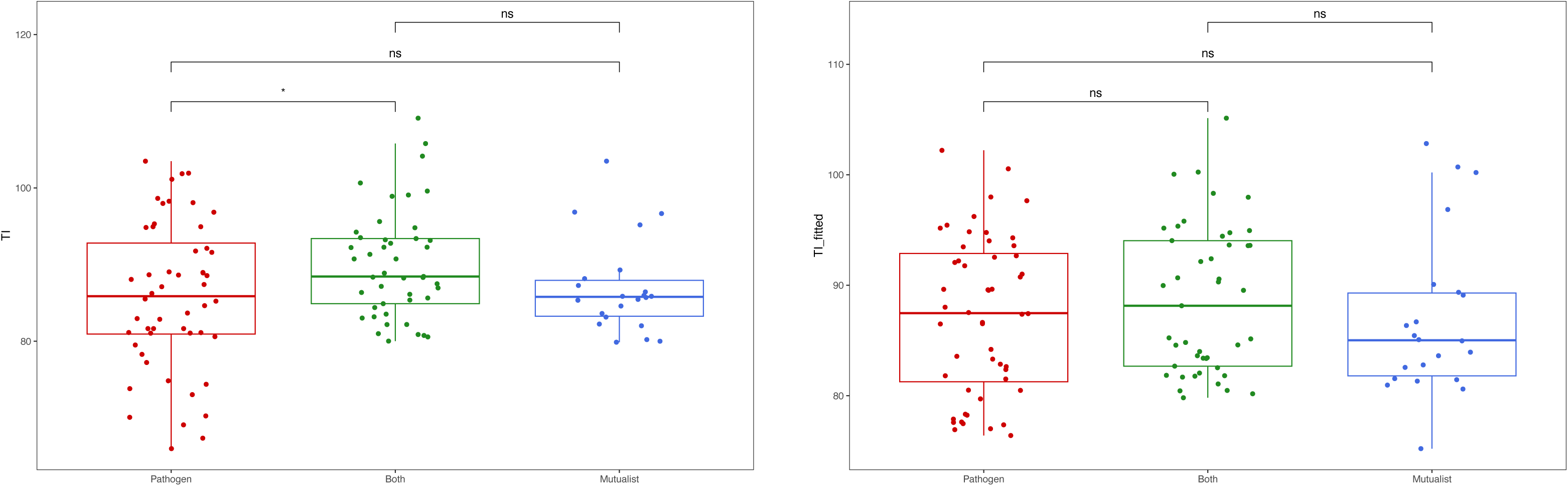
Transient information components against pathogenicity. We show that residuals from a linear model of transient information with genome size separate pathogens from organisms that are both pathogens and mutualists. Significant differences disappear when we plot only the fitted components.

**Supplemental Figure S11.**
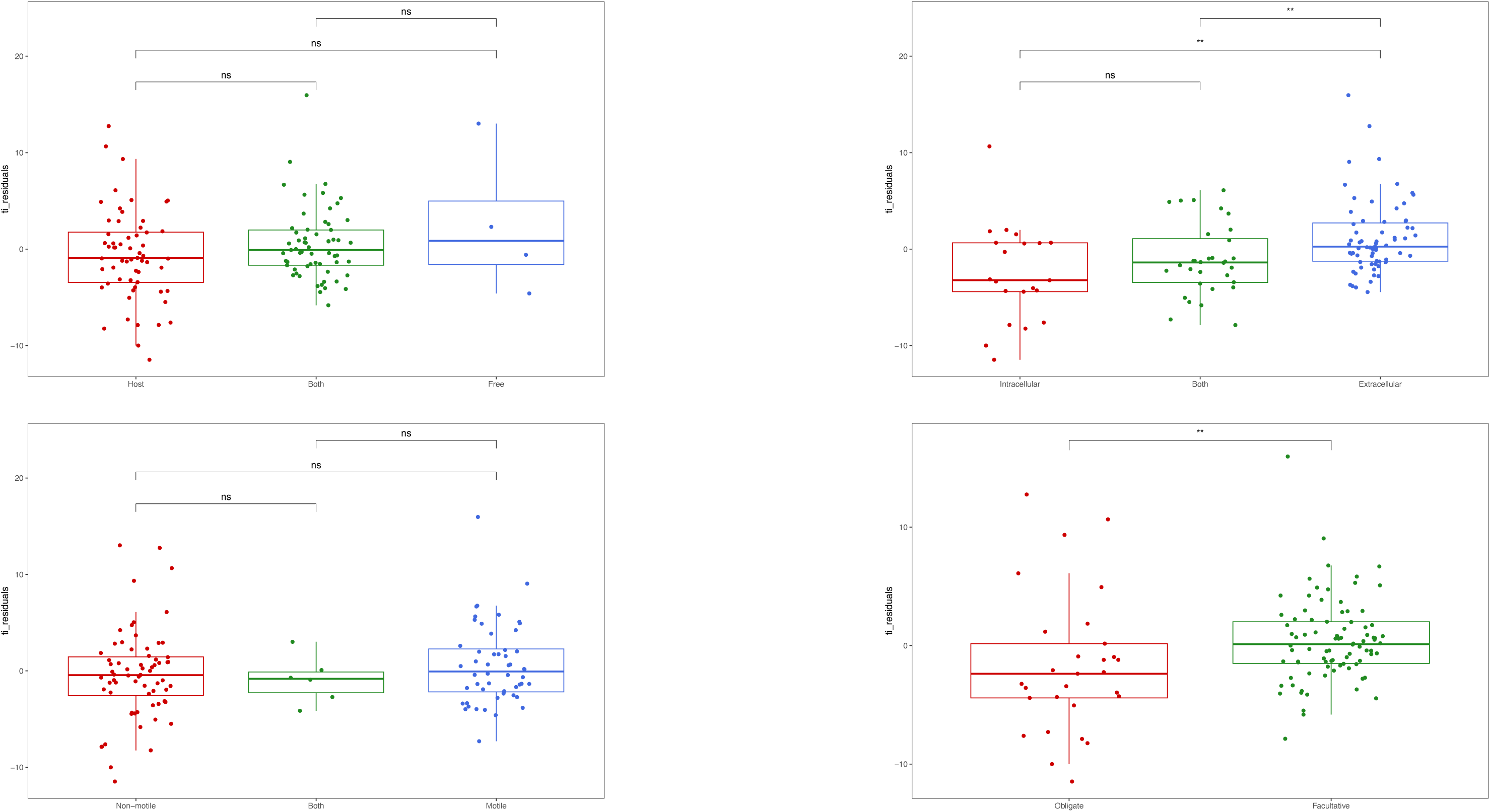
Transient information residuals against four lifestyle traits. We show that transient information residuals can distinguish between organisms that are intracellular versus extracellular, and those that are obligate versus facultative. However, transient information does not yield significant separation between organisms that are pathogens versus non-pathogens, and those that are motile versus non-motile.

## Notes

### Competing Interest Statement

The authors have declared no competing interest.

### Summary of Updates

Methods revised, figures revised, more data and revamped structure.

